# Discovery and characterization of a pigment produced by *Streptococcus pyogenes*

**DOI:** 10.64898/2026.06.05.730339

**Authors:** Sobita Pathak, Theo DeVinney, Artemis Gogos, Reginald A. Woods, Caleb M. Anderson, Jennifer C. Chang, Michael J. Federle

## Abstract

*Streptococcus pyogenes* is a major human-restricted pathogen capable of both localized and systemic diseases, as well as causing post-infectious acute and chronic rheumatological conditions. Despite its well-documented clinical importance and over a century of extensive research, gaps persist in understanding this pathogen. We report the discovery of a novel pigment produced by *S. pyogenes* when cultured in a replete, chemically defined medium. Color development accumulates during growth, requires exposure to oxygen, and remains associated with the bacterial cell. Though only 20% of a small strain collection produced the pigment (8 of 40), positive cultures were overrepresented by M1 and M89 serotypes. Given that pigments are critical virulence and fitness determinants in pathogens like *Staphylococcus aureus and Streptococcus agalactiae*, here we describe initial attempts to characterize *S. pyogenes* pigment biosynthesis, regulation, and potential benefits to the organism. 20,405 transposon mutants were screened for pigment loss in liquid culture, and we identified 94 independent hits enriched in pathways associated with isoprenoid biosynthesis, purine biosynthesis, guanosine transport, and mixed acid fermentation. Although color development requires oxygen, the extracted pigment did not provide antioxidative activity as compared to non-pigmented extracts. Pigment production was inhibited when the Rgg2/Rgg3 quorum-sensing system was active, though by unknown means. Treatment of RAW-Blue macrophages with the extracted pigment significantly reduced NFκB activation, suggesting a potential anti-inflammatory effect. Overall, this study describes a previously uncharacterized pigment produced by *Streptococcus pyogenes* and provides insights into its oxygen-dependent production and associated metabolic pathways.

**SIGNIFICANCE:** *Streptococcus pyogenes* is a ubiquitous pathogen responsible for ∼1.8 million severe infections and 500,000 deaths annually. This organism has not been recognized as a pigment-producing bacterium, but here we identify and characterize the production of a previously unreported colorful compound generated when cultured in a chemically defined medium. This finding reveals an underexplored aspect of *S. pyogenes* biology and raises the question of its contribution to pathogenesis. Given the established roles of microbial pigments in virulence and immune modulation in other pathogens, this work provides a foundation for future investigations into its functional significance and potential as a target for therapeutic or vaccine development.

## INTRODUCTION

*Streptococcus pyogenes*, commonly referred to as Group A streptococcus (GAS), is a Gram-positive, human-restricted bacterium that causes a spectrum of diseases, ranging from acute pharyngitis to skin and soft tissue infections, such as impetigo and cellulitis, to more invasive infections like necrotizing fasciitis and toxic shock syndrome^1,2^. An extensive virulence arsenal governed by a complex regulatory network enables this pathogen to oppose multiple aspects of the host’s immune system, which helps to explain the diverse pathogenicity observed in the clinic ^3–6^. Despite more than a century of study, several *S. pyogenes* virulence-associated factors remain poorly characterized or entirely undocumented, exposing a major gap in our understanding of *S. pyogenes* pathogenesis.

Globally, *S. pyogenes* is responsible for approximately 1.8 million severe infections and 500,000 deaths annually^2^. Beta-lactam antibiotics remain the first-line treatment for infection^7^. However, high rates of antibiotic treatment failure, along with evidence that recent clinical isolates contained rare mutations in the penicillin binding protein 2B, leading to reduced susceptibility to β-lactam antibiotics, underscores an urgent need for alternative therapeutic strategies against *S. pyogenes*^7^. Moreover, no approved vaccine is currently available^8,9^ and prior immunization efforts targeting M protein and Group A carbohydrate (GAC) have been limited by autoimmune complications^9–11^. Together, these limitations highlight critical gaps in our understanding of *S. pyogenes* biology and emphasize the need to identify new virulence-associated pathways that can be therapeutically exploited.

Although well-documented as critical virulence attributes in closely related species, pigmented compounds are not thought to be produced by *S. pyogenes*, as standard culturing conditions yield no colorful products. Pigments are ubiquitous in nature, found in bacteria, fungi, plants, and animals, and are produced as secondary metabolites that, although non-essential for growth, mediate critical interactions with an organism’s environment^12,13^. Light absorption of specific wavelengths defines the colors seen by the eye, but these small molecules perform diverse functions, often protecting cells from stress while also promoting bacterial virulence, as fundamental examples of their ecological and biomedical significance^12,13^. Across multiple clinically relevant bacteria, pigment production has emerged as a conserved strategy to modulate host-pathogen interactions and promote disease progression^14,15^. For example, the human pathogen *Staphylococcus aureus* produces staphyloxanthin, an orange-red triterpenoid carotenoid pigment whose conjugated double-bond system quenches reactive oxygen species and detoxifies free radicals^16,17^. The loss of pigment production renders *S. aureus* vulnerable to destruction by human and murine neutrophils and whole blood^18,19^. Similarly, *Streptococcus agalactiae* (Group B Streptococcus, GBS) produces an orange-red hemolytic pigment called granadaene which contains a polyene of 12 conjugated bonds extended between ornithine and rhamnose^20,21^. Granadaene, like staphyloxanthin, has antioxidant properties and enhances GBS survival within macrophages^22,23^. Pigmented GBS exhibits a higher survival rate in a systemic murine infection model compared to non-pigmented GBS^22^. *Pseudomonas aeruginosa* produces pyocyanin, a blue-green pigment derived from phenazine^15^. In contrast to the antioxidant properties of staphyloxanthin and granadaene, pyocyanin displays a paradoxical pro-oxidant activity. It accepts electrons from cellular reducing agents, such as NADPH and reduced glutathione, and subsequently transfers them to molecular oxygen, generating reactive oxygen species (ROS), including hydrogen peroxide and singlet oxygen^24^. This process occurs at the expense of host antioxidant defenses, particularly glutathione and catalase^24,25^. Generated ROS exert antibacterial activities against competing organisms like staphylococci, streptococci, pneumococci, *V. cholerae,* and gonococci^15^. These examples highlight pigment production as virulence strategies in both Gram-positive and Gram-negative pathogens, by modulating redox balance, enhancing survival in host environments, and mediating inter-microbial competition.

Here, we describe the identification of a pigment produced by *S. pyogenes* under specific growth conditions. We identify genes associated with pigment biosynthesis and regulation, and we identify means to extract and purify the compound to assist in the elucidation of its biological activities. Given the well-established roles of pigments in the virulence and fitness of other bacterial pathogens, we hypothesize that this novel pigment contributes to *S. pyogenes* pathogenicity and may offer a novel therapeutic or vaccine target. Accordingly, in this study, we aimed to (*i*) define the conditions that promote pigment production, (*ii*) elucidate its biosynthetic pathway using transposon mutagenesis, and (*iii*) assess its biological relevance.

## RESULTS

### *S. pyogenes* cultured in CDM produces an uncharacterized pigment

Pigment production has not been documented in *S. pyogenes*^26^. We found that cell pellets collected after culturing in a peptide-free, chemically defined, nutritionally replete medium (CDM)^27^contained an uncharacterized pigment (Figure 1A). No pigmentation was observed in cell pellets after static growth in the complex medium Todd-Hewitt with yeast extract (THY). The color of the pigment intensified once cells were exposed to air, with robust development occurring if cultures were aerated vigorously prior to pelleting (Figure 1B). Pigment production was not visible in colonies of bacteria grown on CDM-based solid agar.

**Figure 1.**
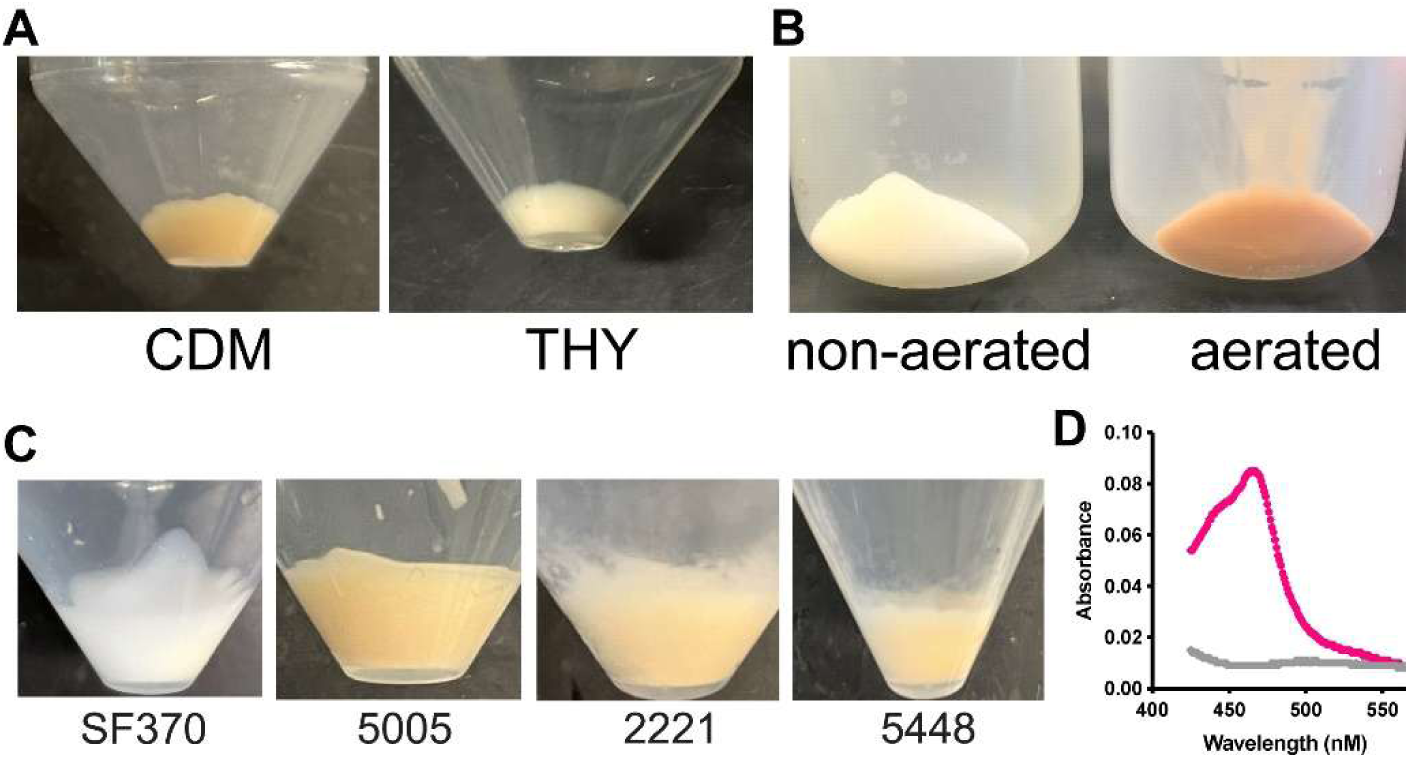
*S. pyogenes* produces a pink-orange pigment when grown in a chemically defined medium (CDM). **(A)** Overnight culture pellets of MGAS5005 grown in CDM or Todd Hewitt yeast (THY) media. **(B)** Aeration-dependent intensification of visible pigment. **(C)** Representative pigment formation in different M1 strains. **(D)** UV–visible spectra of pellet extracts from pigmented (MGAS5005) and non-pigmented (NZ131) strains.

Pigment production was strain and serotype dependent. Of 42 strains tested, consisting of 23 different serotypes, production was observed in 3 of 4 serotype-M1 strains (MGAS5448, 2221, and 5005, but not SF370), 2 of 6 M89 strains (MGAS2017 and 58362), and in only 3 other strains of other serotypes (M96, M101, st980584) (Supplementary Table 3). Representative images of pigment formation in M1 strains are shown in Figure 1C. Among these strains, MGAS5005 generated the greatest amount of color development and therefore was used for most experiments henceforth. To the eye, the pigment color appeared as pink-orange, consistent with a maximal light absorbance observed at 468 nm (Figure 1D).

### Role of oxygen in color development

We observed that exposure of MGAS5005 cultures to air resulted in a marked intensification of pellet coloration, suggesting a role for oxygen in pigment formation. To directly assess oxygen dependency, cultures were first grown under strictly anaerobic conditions using an anaerobic chamber. Cells cultivated anaerobically produced uncolored pellets. However, upon subsequent exposure to ambient air, these pellets rapidly developed a pink coloration within approximately 5 minutes (Figure 2A). Cultures grown statically (non-shaking) under normal atmospheric conditions supplemented with 5% CO_2_ formed visibly pink pellets during growth, and further aeration (shaking) of stationary-phase cultures intensified the color toward orange (Figure 2A). Consistent with a requirement for oxygen, addition of the oxygen-scavenging reagent Oxyrase to CDM prior to microaerophilic culturing prevented color development (Figure 2B). However, neither the addition of Oxyrase to already pigmented cultures nor the placement of pigmented bacteria into an anaerobic chamber reverted them to a colorless state. Conversely, addition of hydrogen peroxide (H_2_O_2_), which can generate hydroxyl radicals, through Fenton chemistry^28^, to stationary-phase cultures, increased pigment intensity in a concentration dependent manner resulting in a visibly darker pigment pellet (Figure 2C). Treatment of anaerobically cultured cells with chloramphenicol, a protein translation inhibitor, immediately prior to oxygen exposure developed color indistinguishable from that of drug-treated and untreated cells under both aerobic and anaerobic conditions (Figure 2D). These results indicated that *de novo* protein synthesis was not required for color formation upon exposure to oxygen and that compound(s) comprising the pigment must have been produced prior to oxygen exposure. Moreover, these results indicated that oxygen-related stress was not required for pigment production.

**Figure 2.**
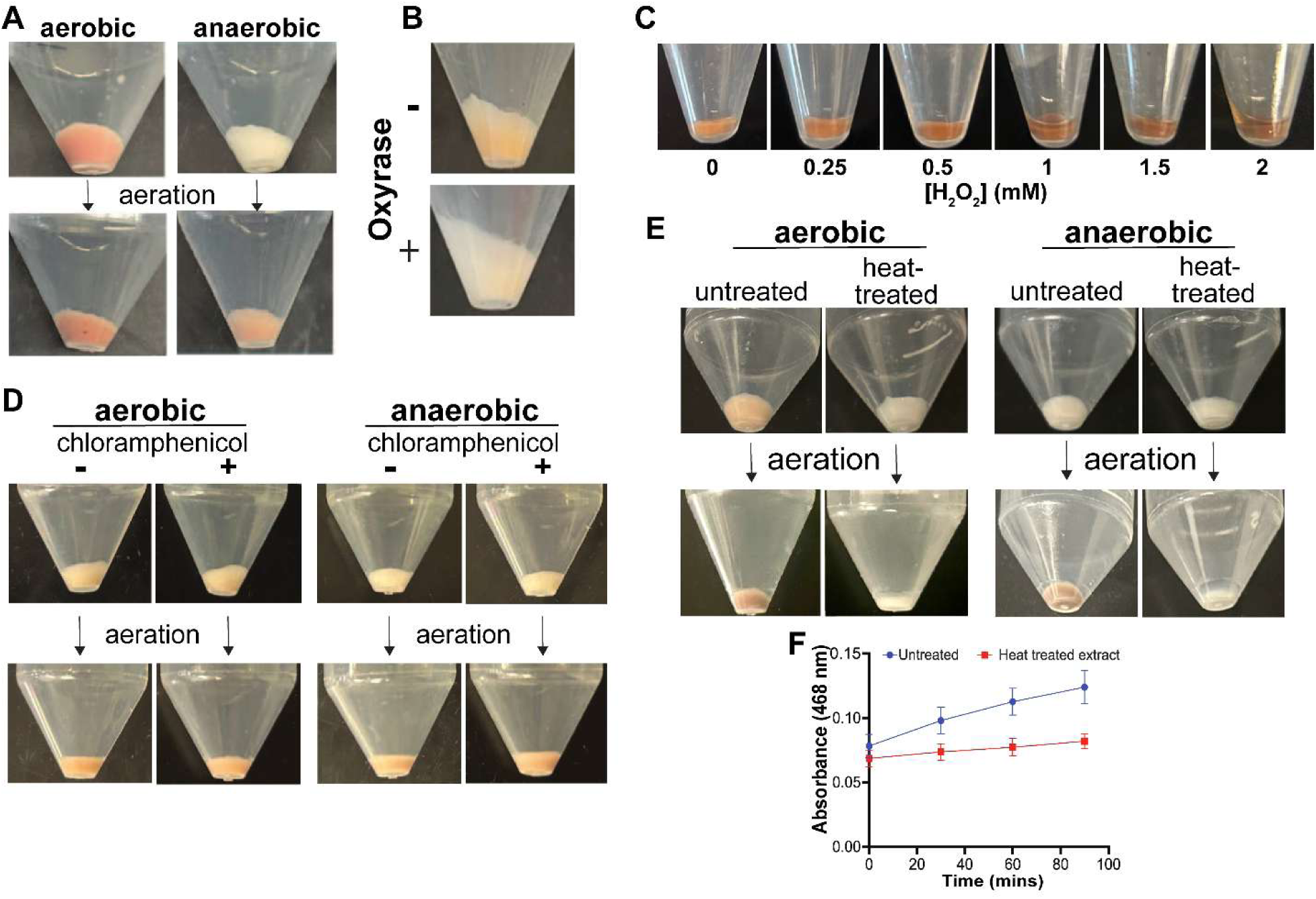
Oxygen is required for color development. **(A)** Cell pellets of MGAS5005 grown aerobically are pigmented, whereas pellets from anaerobically grown cultures are colorless until exposure to air. **(B)** Addition of oxyrase prevented the production of pigment. **(C)** Addition of H₂O₂ intensified the pigment color. **(D)** Cell pellets treated with chloramphenicol (3 µg/mL) prior to aeration maintained the ability to develop color. **(E)** Pellets from heat-inactivated cultures did not develop color. **(F)** Methanol extracts from anaerobically grown cultures developed color upon exposure to air, whereas heat-treated samples showed delayed color development.

To determine whether color development required an enzymatic catalyst, cultures grown both aerobically and anaerobically were heat-treated at 80°C for 30 mins prior to centrifugation. Heat-inactivated cultures failed to develop color even after prolonged exposure to air (Figure 2E), indicating that color formation could not occur in metabolically inactive cells. Heat treatment of the developed pigment did not cause a loss of color, ruling out the possibility that the colored compound itself was susceptible to heat (Figure S1). Anaerobically cultured cell pellets, extracted with methanol under anaerobic conditions and subsequently exposed to air, retained the ability to develop color, albeit slowly (Figure 2F). Anaerobic extracts subjected to heat decreased but did not eliminate modest color development. Collectively, these results show that oxygen is required for a heat-labile, color-development reaction.

### Transposon mutagenesis reveals pathways important for pigment production

To identify genes that contribute to pigment production, the plasmid-based transposon IS*S1* was utilized to generate insertion mutations in MGAS5005^29^. As bacterial colonies grown on agar plates did not produce enough pigment to be seen by eye, isolated mutant colonies were picked into 0.2 ml CDM in 96-well plates that were sealed with a gas-permeable film and incubated overnight. Cultures were subsequently pelleted by centrifugation and screened by eye for the absence of pigmentation. 20,405 mutant clones were screened, identifying 94 non-pigmented clones that were validated as being colorless after subsequent culturing in 5 ml of CDM + erm (1 µg/mL). The genomic location of transposon insertion sites for each clone was determined after isolation of chromosomal DNA and sequencing of transposon junctions^29,30^. Functional annotation revealed that several mutants could be assigned to four major metabolic pathways: mixed acid fermentation, purine biosynthesis, guanosine transport, and the isoprenoid pathway (Table 1, Figure 3).

**Figure 3:**
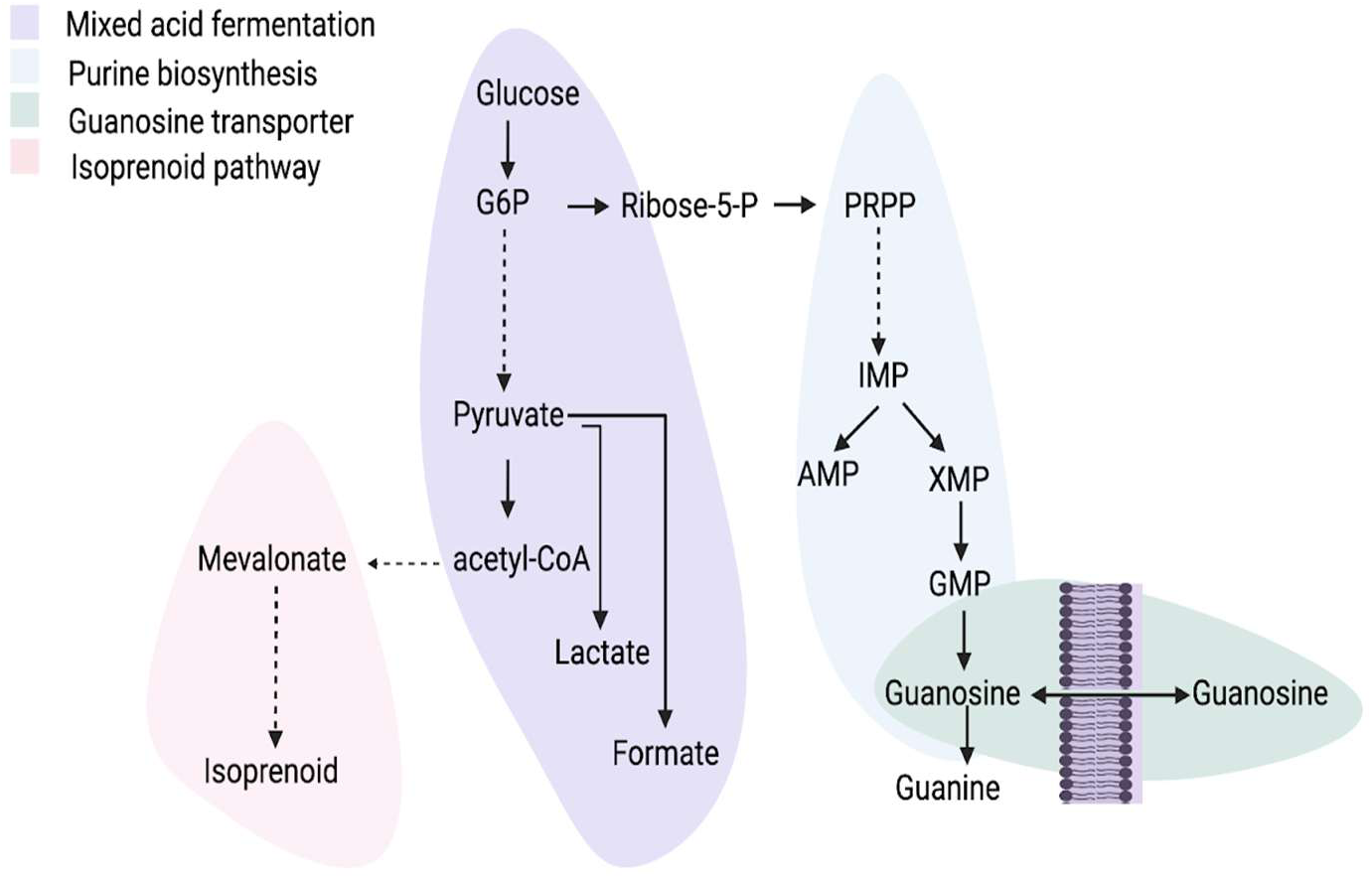
Proposed model illustrating the integration of purine metabolism, guanosine transport, and isoprenoid biosynthesis in pigment biosynthesis.

**Table 1.** IS*S1* insertion mutants associated with loss of pigmentation.

|  | **Spy designation** | **No of hits** | **Gene name** | **Description** |
| --- | --- | --- | --- | --- |
| **Mixed Acid Fermentation, Isoprenoid pathway** | *M5005_spy1569* | 7 | *pfl* | Formate acetyltransferase |
|  | *M5005_spy1238* | 3 | *artQ* | Arginine transport system permease protein (*fps* is downstream (*M5005_spy1231*)) |
|  | *M5005_spy1344* | 1 | *atoB* | Acetoacetyl CoA |
|  |  | **Total hits = 11** |  |  |
| **Purine biosynthesis** | *M5005_spy0022* | 2 | *purC* | SAICAR synthetase, purine biosynthesis |
|  | *M5005_spy0023* | 4 | *purL* | FGAM synthetase, purine biosynthesis |
|  | *M5005_spy0024* | 3 | *purF* | Amidophosphoribosyltransferase , purine biosynthesis |
|  | *M5005_spy0025* | 2 | *purM* | AIR synthetase, purine biosynthesis |
|  | *M5005_spy0026* | 3 | *purN* | GAR transformylase, purine biosynthesis |
|  | *M5005_spy0029* | 1 | *purD* | Phosphoribosylamine-glycine ligase |
|  |  | **Total hits = 15** |  |  |
| **Nucleoside transport** | *M5005_spy0939* | 3 | *nupQ* | Nucleoside transport system permease protein |
|  | *M5005_spy0940* | 3 | *nupP* | Nucleoside transport system permease protein |
|  | *M5005_spy0941* | 3 | *nupO* | Nucleoside transport ATP binding protein |
|  | *M5005_spy0942* | 1 | *nupN* | Nucleoside binding protein |
|  | *M5005_spy0943* | 1 | *cdd* | Cytidine deaminase |
|  |  | **Total hits = 11** |  |  |
| **Rgg2/3 QS-related** | *M5005_spy0367* | 2 | *mtsR* | Metal dependent transcriptional regulator |
|  | *M5005_spy0369* | 1 | *mtsC* | Manganese transport system membrane protein |
|  | *M5005_spy0440* | 1 | *rgg3* | Transcriptional repressor |
|  | *M5005_spy1782* | 1 | *pepO* | Neutral endopeptidase |
|  |  | **Total hits = 5** |  |  |
| **Other transcriptional regulators** | *M5005_spy0122* | 3 | *serR/ yheO* | Putative DNA binding protein, YheO transcriptional regulator |
|  | *M5005_spy0282* | 1 | *covR* | CovR response regulator |
|  | *M5005_spy0947* | 3 | *ciaH* | Sensor protein, *ciaHR* two component system |
|  | *M5005_spy1720* | 1 | *mga* | Transacting positive regulator |
|  |  | **Total hits= 6** |  |  |
| **Riboflavin biosynthesis** | *M5005_spy0496* | 4 | *yigB* | Haloacid dehydrogenase (HAD) superfamily, FMN and 5-amino-6-(5-phospho-D-ribitylamino) uracil phosphatase |
| **Zinc metabolism** | *M5005_spy0077* | 2 | *adcR* | Transcriptional regulator, MarR family |
|  | *M5005_spy0079* | 1 | *adcB* | High-affinity zinc uptake system membrane protein |
| **Methionine transport** | *M5005_spy0271* | 1 | *atmB* | ABC transporter substrate-binding protein, *nlpA* |
|  | *M5005_spy0272* | 1 | *atmD* | ABC transporter ATP binding protein, also called *abcC* |
| **Multiple hits** | *M5005_spy0320* | 2 |  | Hypothetical cytosolic protein |
|  | *M5005_spy0607* | 3 | *gacF* | GAC biosynthesis |
|  | *M5005_spy0794* | 2 | *thdF* | GTPase |
|  | *M5005_spy1837* | 2 | *gdpP* | c-di-AMP phosphodiesterase, GGDEF domain |
| **Single hits** | *M5005_spy0087* | 1 | *comYB* | ABC transporter subunit |
|  | *M5005_spy0093* | 1 |  | adenine-specific methyltransferase |
|  | *M5005_spy0114* | 1 | *srtA* | Class c sortase |
|  | *M5005_spy0247* | 1 |  | Serine-type D-Ala-D-Ala carboxypeptidase |
|  | *M5005_spy0320* | 1 |  | Putative cytosolic protein |
|  | *M5005_spy0332* | 1 |  | Unannotated gene |
|  | *M5005_spy0477* | 1 |  | Putative membrane spanning protein |
|  | *M5005_spy0487* | 1 |  | Putative exported protein |
|  | *M5005_spy0545* | 1 | *agaS* | Galactosamine-6-phosphate deaminase (isomerizing) |
|  | *M5005_spy0547* | 1 |  | Phosphoesterase |
|  | *M5005_spy0561* | 1 | *epf* | Putative extracellular matrix-binding protein, cell wall anchor |
|  | *M5005_spy1085* | 1 | *bglA* | Beta-glucosidase |
|  | *M5005_spy1104* | 1 |  | cytosolic protein |
|  | *M5005_spy1121* | 1 | *ptsH* | Phosphocarrier protein HPr |
|  | *M5005_spy1176* | 1 |  | Phage protein |
|  | *M5005_spy1215* | 1 |  | Unannotated gene |
|  | *M5005_spy1339* | 1 | *priA* | Primosomal protein N |
|  | *M5005_spy1481* | 1 | *manN* | PTS system, mannose-specific IID component |
|  | *M5005_spy1529* | 1 | *shp* | Putative heme binding protein |
|  | *M5005_spy1571* | 1 |  | C3-degrading proteinase |
|  | *M5005_spy1603* | 1 |  | Putative alkaline-shock protein |
|  | *M5005_spy1783* | 1 | *dexS* | Trehalose-6-phosphate hydrolase |
|  | *M5005_spy1794* | 1 |  | Membrane spanning protein |
|  | *M5005_spy1799* | 1 | *recA* | Recombination protein |
|  | *M5005_spy1823* | 1 |  | Integral membrane protein |

### Role of isoprenoid biosynthesis in pigment production

The identification of mutations in genes *pfl*, *atoB*, and *artQ* suggested a connection between the mixed acid fermentation pathway with mevalonate biosynthesis (Figure 4A). The mevalonate pathway produces precursors for isoprenoid and carotenoid compounds that account for many types of pigments. The spectrophotometric profile of the GAS pigment (max absorbance 468 nm) is consistent with carotenoid-containing pigments like staphyloxanthin (max 478 nm) and granadaene (455–460 nm)^16,20^.

**Figure 4:**
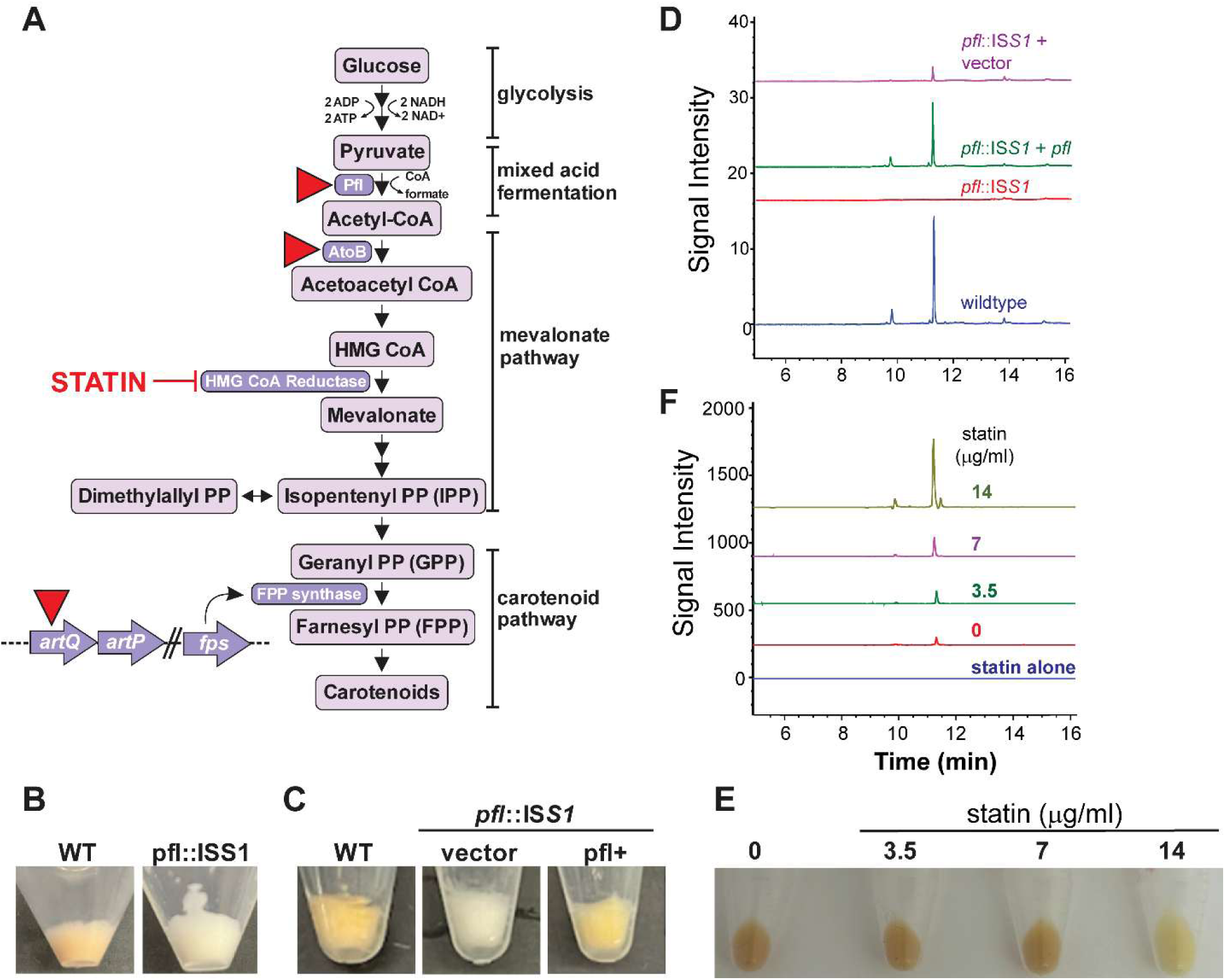
The isoprenoid biosynthetic pathway is important for pigment production. **(A)** Schematic representation of the isoprenoid biosynthetic pathway indicates genes identified as hits (red triangles) in the transposon mutagenesis screen. **(B)** Visualization of color differences between WT and *pfl*::IS*S1* mutant. **(C)** Complementation of *pfl* restored the pigmented phenotype. **(D)** HPLC analysis of extracts obtained from WT, *pfl*::IS*S1* and *pfl*-complemented strains. **(E)** Simvastatin treatment altered pigment color from orange to yellow at subinhibitory concentrations. **(F)** Simvastatin treatment enhanced the 468nm absorbance peak intensity.

The pyruvate formate lyase enzyme (Pfl) provides one of two parallel mixed acid fermentative pathways that catalyze the conversion of pyruvate to acetyl-CoA, the other utilizes the pyruvate dehydrogenase complex (Pdh). Pfl is a glycyl radical-containing enzyme, sensitive to oxygen and therefore active only under anaerobic or microaerophilic conditions^31^. Seven independent insertions in the *pfl* gene were isolated, the highest number among all identified genes. Loss of pigmentation in a *pfl*::IS*S1* mutant was confirmed visually (Figure 4B) and no absorbance peak was detected at 468 nm in the UV–Vis spectrum (Figure S2). Methanol extracts of pelleted cells, evaluated by reverse-phase HPLC at an absorbance of 468 nm displayed differences between the WT and *pfl::ISS1* mutant. Two peaks were observed at 10.017 min and 11.435 min for WT that was absent in the *pfl::ISS1* extract (Figure 4D). Complementation of the *pfl::ISS1* mutant with *pfl* expressed in trans restored pigment production, as confirmed both visually and by HPLC analysis (Figure 4C, D). An in-frame deletion of *pfl* was also generated and complemented, with loss and restoration of pigment, respectively (Supplementary Figure S3).

The product of Pfl, acetyl-CoA, can then be converted to acetoacetyl-CoA by thiolase AtoB (1 hit), providing substrate for the mevalonate pathway. Three mutants of *artQ* were identified, and while this gene and the downstream-adjacent *artP* encode a putative ABC-type arginine transporter, we considered the possibility of polar effects by the insertions could affect expression of *fps*, which is the seventh gene downstream of *artQ* in a likely polycistronic operon. Farnesyl pyrophosphate synthase (FPS) is a key enzyme in the carotenoid pathway that converts geranyl pyrophosphate (GPP) to farnesyl pyrophosphate (FPP)^32^. Thus, we reasoned that pigment production could depend on mevalonate for the incorporation of an isoprenyl or carotenoid component. We therefore generated a chromosomal in-frame deletion of *fps* and trans-complementing plasmid. Effects of the deletion were minor but noticeable, and complementation boosted pigmentation (Figure S3); however, deleting *fps* did not phenocopy the *artQ*::IS*S1* mutant and therefore its effect on pigment is unclear.

To test the possibility that mevalonate contributes a key precursor to pigment biosynthesis, simvastatin, a well-known inhibitor of the rate-limiting enzyme HMG-CoA reductase that converts HMG-CoA to mevalonate^33^ was applied to cultures. As HMG-CoA reductase is critical in isoprenoid production and viability^34^, a titration of concentrations of simvastatin allowing growth were tested, identifying 15 µg/mL as the minimal inhibitory concentration (MIC). A noticeable change in pigment intensity and hue was observed as concentrations approached the MIC (Figure 4E, F). Unexpectedly, rather than eliminating pigment production, simvastatin-treated cultures changed the pigment color from orange to yellow and extracts of these cells analyzed by HPLC displayed an increased signal at 468 nm. This result confirmed that disrupting the mevalonate pathway changed the pigment’s characteristics, but it remains unclear whether it is due to depleting IPP availability (the product of the mevalonate pathway) or the effects of the possible buildup of HMG-CoA.

### Purine biosynthesis is important to pigment production

In *S. pyogenes*, nucleotide pools are generated and levels balanced by *de novo* synthesis and salvage pathways. Our screen identified nonpigmented mutants with IS*S1* insertions in purine biosynthesis (15 hits across 6 genes) and nucleoside salvage pathways (11 hits across 5 genes). Although GAS contains all necessary purine biosynthetic genes needed to produce adenine and guanine nucleotides, CDM is replete with these purines (as well as the pyrimidine uracil). It was therefore surprising that *pur* gene mutations impacted pigment development when the bacteria were able to scavenge bases. To confirm the importance of the *de novo* pathway, we used mycophenolic acid (MPA), a noncompetitive, selective, and reversible inhibitor of inosine monophosphate dehydrogenase (IMPDH, encoded by *guaB*) that converts inosine monophosphate (IMP) to xanthosine 5’ monophosphate (XMP)^35^ (Figure 5A). MPA treatment eliminated pigment production in cell pellets and extracts (Figure 5B, C).

**Figure 5:**
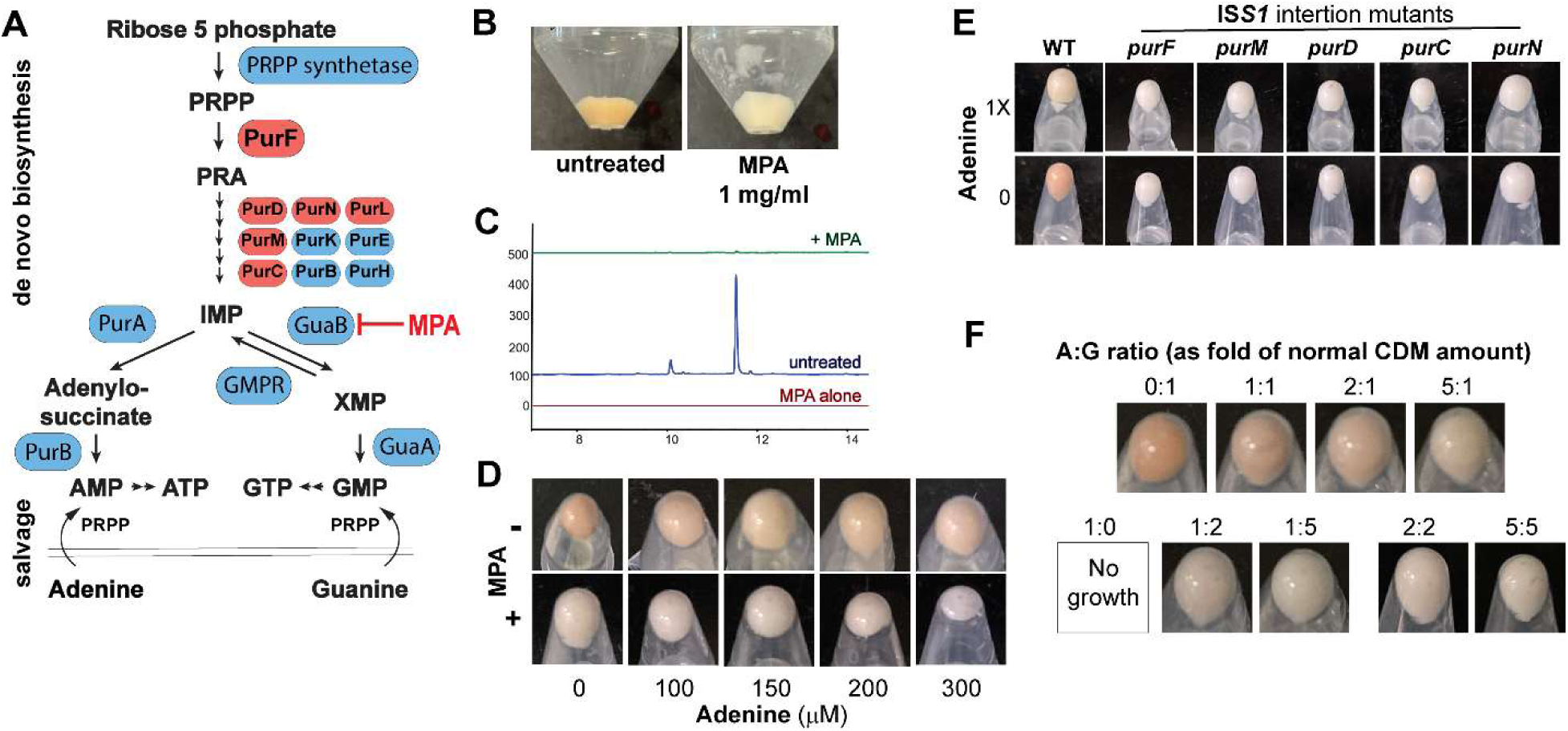
The impact of purine metabolism on pigment production. **(A)** *S. pyogenes* purine *de novo* biosynthesis and salvage pathways. Enzymes bordered in red indicate genes that were disrupted by IS*S1*. **(B)** Treatment with 1 mg/ml mycophenolic acid (MPA), an inhibitor of GuaB, prevented pigment production. **(C)** HPLC chromatograms comparing MPA-treated and untreated culture extracts (red line shows absorbance profile of MPA alone in solution). **(D)** Removal of adenine from CDM results in hyperpigmentation, and the effect of MPA. **(E)** Pigmentation of *pur* mutants grown with and without adenine. **(F)** Effects of adenine: guanine ratios in growth medium. Amounts of adenine (150 µM) and guanine (132 µM) are shown as fold amounts present in CDM.

Because MPA blocks the conversion of IMP to XMP (and therefore inhibits guanosine nucleotide production) but does not block production of succinyl-adenosine monophosphate (S-AMP), we hypothesized that disruption in nucleotide-pool balance may be responsible for the pigment inhibition caused by MPA. We then tested whether altering the ratios of exogenous purines would change pigment production and found that removing adenine resulted in a robust increase in pigment (Figure 5D). Although restricting adenine in cultures of *pur* mutants did not restore pigmentation, (Figure 5E), adenine depletion did overcome the inhibitory effect of MPA (Figure 5F). This supports the notion that buildup of adenosine nucleotides caused by MPA could be overcome by limiting the availability of adenine provided exogenously.

To explore further the role of purines, we varied the concentrations of guanine and its ratio to adenine. Excess guanine led to a dose-dependent decrease in pigmentation, with higher concentrations ultimately abolishing pigment entirely (Figure 5G). Together, these results indicated that pigment production in *S. pyogenes* is highly sensitive to changes in adenine and guanine ratios and suggest that well-known substrate inhibition mechanisms resulting in negative feedback on purine metabolic pathways may account for pigment yield. Furthermore, GTP serves as a precursor for riboflavin biosynthesis. Although GAS does not appear to possess a complete riboflavin biosynthetic pathway, it is notable that *yigB* was hit 4 times in transposon mutagenesis and a generated clean in-frame deletion showed a strong, complementable phenotype (Supplementary Figure S3). This observation further prompted us to examine whether GAS pigment shared characteristics with riboflavin. Comparison of HPLC retention times and UV visible absorption profiles, however, indicated that the GAS pigment is distinct from riboflavin (Supplementary Figure S4).

### Activation of Rgg2/Rgg3 QS inhibits pigment production

Non-pigmented transposon mutants were identified in *rgg3*, *pepO*, and *mtsR*, all genes that we previously described as affecting the Rgg2/Rgg3 quorum sensing (QS) system. This QS system involves two transcriptional regulators, Rgg2 (activator) and Rgg3 (repressor), which are modulated by SHP pheromones (short hydrophobic peptides). Deletion of *rgg3* eliminates repression on the QS system and causes the up-regulation of the pheromones and QS-regulated genes (e.g., *stcA* and *qimJ*)^36–38^. The promoter of *shp3* is also governed by MtsR, a metalloregulator that represses *shp* expression. PepO is an endopeptidase that inactivates SHP signaling by proteolyzing the pheromones. Disruption of any of these genes increases the activation of the QS system, and therefore we interpreted the loss of pigment by IS*S1* insertion in these genes to be an effect of QS activation. We tested this hypothesis by treating MGAS5005 with 100 nM SHP and confirmed the loss of pigment in this culture, as well as for wildtype strains 2221 and 5448 (Figures 6A, 6B). The production of pigment was also tested for in-frame deletions of *rgg2*, *rgg3*, *pepO*, and the QS-regulated *stcA* and *qimJ* mutants. Surprisingly, the Δ*rgg3* mutant maintained its ability to produce the pigment unless exogenous SHP was added, suggesting the loss of Rgg3 in MGAS5005 does not fully induce the system. The double-deletion of *rgg2* and *rgg3*, which lacks both SHP receptors and therefore does not respond to SHP did not decrease pigment production when treated. As predicted, the Δ*pepO* mutant remained in a QS-ON state and blocked pigment production, whereas inhibition of pigment production when QS was active did not depend on *stcA* or *qimJ*.

**Figure 6.**
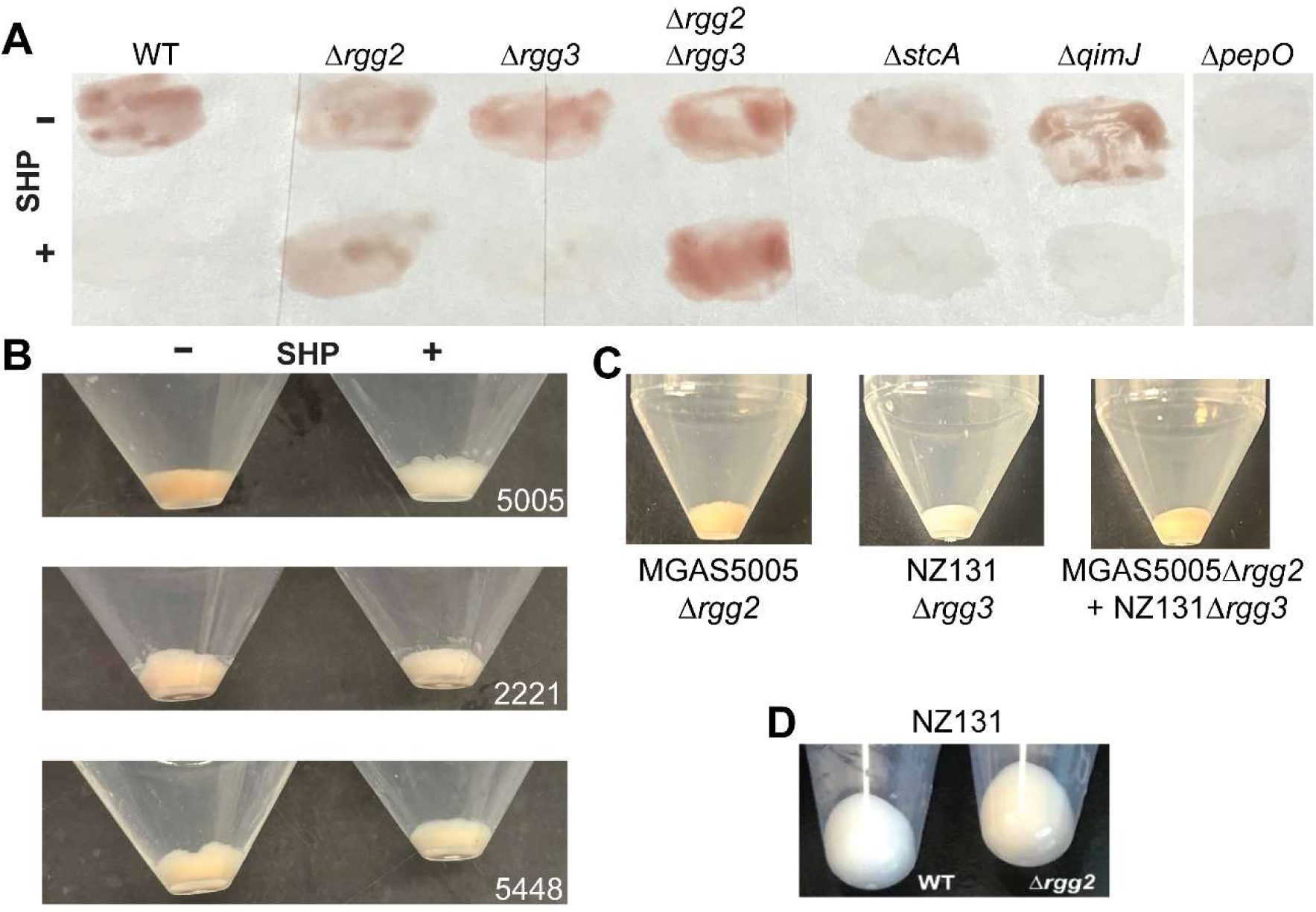
Rgg2/Rgg3 signaling affects pigment production. **(A)** Culture pellets of MGAS5005 mutants, grown without (-) or with (+) 100 nM SHP, were suspended in CDM and spread onto white paper for visualization. **(B)** Strains MGAS5005, 2221, and 5448 were grown without or with 100 mM SHP and pelleted. **(C)** Cultures of MGAS5005 Δ*rgg2* and NZ131 Δ*rgg3* were grown separately or together prior to centrifugation. **(D)** NZ131 Δ*rgg2* mutant produces a small amount of pigment.

We also tested *rgg* mutants in NZ131, which we found to be an otherwise non-pigmented strain in CDM. However, deleting the *rgg2* gene, which prevents activation of the QS system, caused this strain to produce a small but visible amount of pigment (Figure 6D). Co-culturing MGAS5005Δ*rgg2* (QS-OFF) with NZ131Δ*rgg3* (QS-ON) bacteria did not prevent MGAS5005 from producing pigment, discrediting a scenario in which an extracellular product of *S. pyogenes* that is made when QS is active was degrading the pigment (Figure 6C). These results indicate that QS regulates pigment production at the level of biosynthesis rather than pigment stability.

### Pigment extract bioactivity

Pigments can have a variety of biological functions, including providing protection from environmental stresses as free radical scavengers, having antimicrobial and hemolytic activities, and increasing membrane stabilization^39,40^. There are also instances where bacterial pigments modulate host immune responses. Pyocyanin produced by *Pseudomonas aeruginosa* alters the NFκB signaling and cytokine production through redox dependent mechanisms, while the carotenoid pigment staphyloxanthin protects from oxidative burst and contributes to immune evasion^15,19^. Based on this, we tested whether the *S. pyogenes* pigment could modulate an NFκB-based reporter assay, as activation of this pathway is a hallmark of inflammatory responses^41^. To this end, we assessed whether extracts derived from pigmented MGAS5005 and non-pigmented mutant cultures, when applied to cultured macrophages, affected signaling. We employed RAW-Blue macrophages, a reporter cell line engineered to express a secreted embryonic alkaline phosphatase (SEAP) gene under the control of NFκB responsive elements. Bacterial extracts were applied onto these reporter cells in combination with the TLR2/TLR1 agonist Pam3CSK4 to assess whether the pigment exhibits pro- or anti-stimulatory effects on innate immune signaling. Treatment with the pigmented extract resulted in a reduced reporter signal compared to the non-pigmented extract, indicating its ability to inhibit NFκB activation (Figure 7A). We refined the extract further using size-exclusion chromatography and collected colored fractions (Figure S5). A total of 90 colored samples were collected and pooled based on color and spectral absorbance profiles, yielding five major fraction groups: slightly colored (F1), dark orange (F2), orange (F3 and F4), and yellow (F5). Among these, the dark orange fraction (F2) retained inhibitory activity in the NFκB assay (Figure 7B), suggesting that the immunomodulatory activity is associated with a specific component enriched within this fraction. To determine whether the observed change in NFκB signaling was associated with a downstream inflammatory response, nitrite production was measured using a Griess assay in cultured RAW267.4 cells. Consistent with the NFκB assays, unpigmented extracts led to higher nitrite levels than pigmented extracts; however, the magnitude of the effect was modest and varied across strain backgrounds, with only minor differences observed between WT and *pfl-*mutant extracts under the conditions tested. (Figure S6)

**Figure 7:**
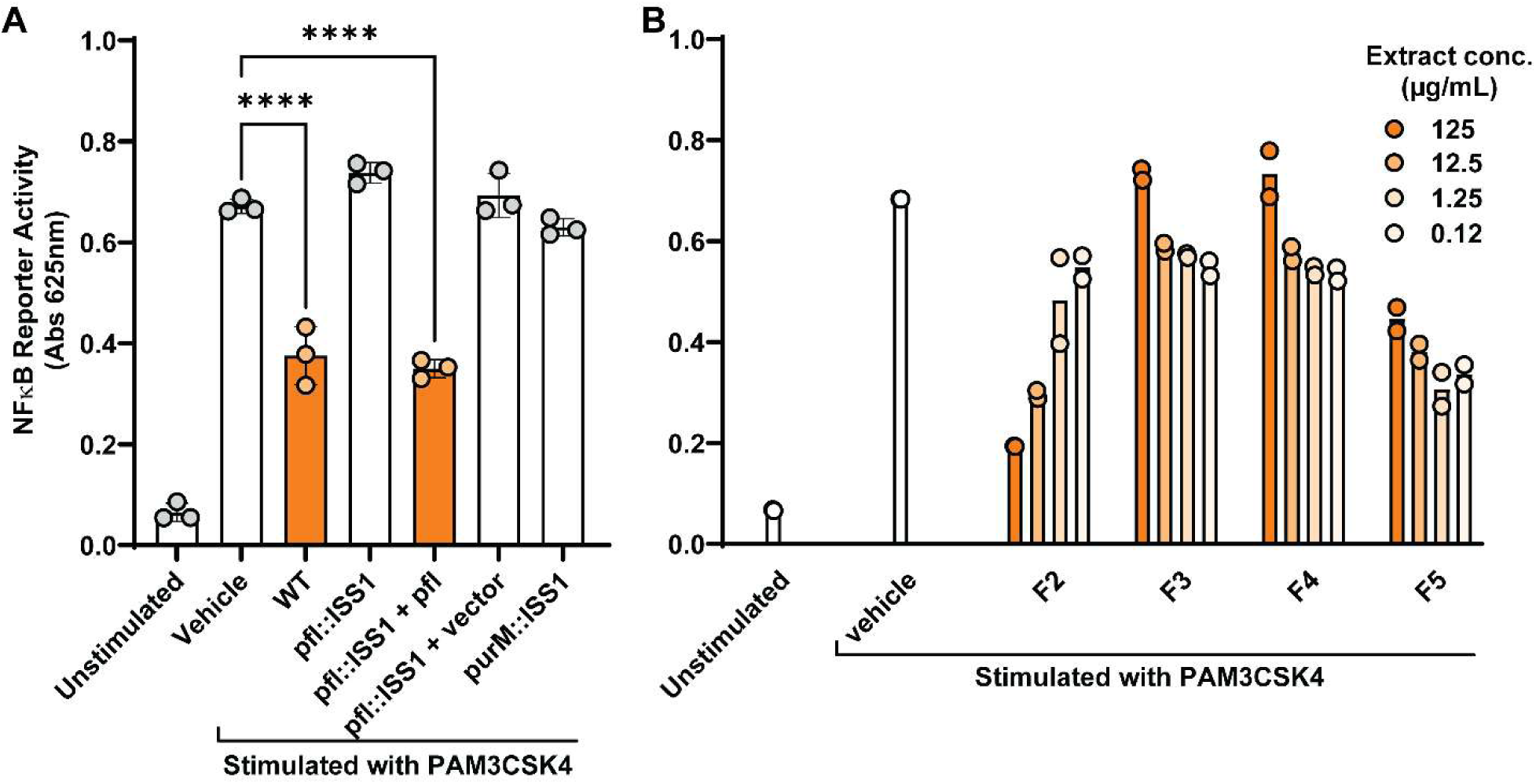
Pigmented extract inhibits the NFκB activation in PAM3CSK4-stimulated RAW-BLUE cells. **(A)** RAW-Blue macrophages were treated with 125 µg/ml of crude extracts derived from indicated strains. Colored bars indicate extracts that appeared pigmented. NFκB activity was quantified by measuring color absorbance (625 nm) of the QuantiBlue reagent due to the secreted alkaline phosphatase reporter. Data are shown as means ± SD from three independent experiments (n = 3). Statistical significance was determined by two-way ANOVA with Tukey’s multiple-comparisons test. ****, *p* < 0.0001. **(B)** NFκB activation was further examined in fractions of WT pigment extracts purified by size-exclusion chromatography, dried, weighed and suspended in PBS (axis values are same as in panel A). Bars indicate the means of two replicate experiments.

We also determined whether the *S. pyogenes* pigment exhibited characteristics seen for pigments of other organisms. Given the dependency of color development on exposure to oxygen, we tested the antioxidant capacity of extracts obtained from pigmented WT and non-pigmented mutant cultures using the colorimetric 2,2′-azinobis-3-ethylbenzothiazoline-6-sulfonic acid (ABTS) assay. However, no significant differences were observed between the two types of extracts (Figure S7). Antibacterial activity of the extracts was tested by disc-diffusion assays against *Pseudomonas aeruginosa*, *Staphylococcus aureus*, *Streptococcus agalactiae*, *Escherichia coli*, and *S. pyogenes* strains MGAS5005 and NZ131. No zones of inhibition were observed for any of the organisms when 10 µg of extract was applied (Figure S8). Extracts spotted onto blood agar plates did not lead to the lysis of red blood cells and when live MGAS5005 (WT) and non-pigmented mutants were stabbed into the agar, which provides a means to assess hemolytic activity due to secreted streptolysins O and S, no differences in hemolytic zones were noticeable. (Figure S9).

## DISCUSSION

Environmental conditions are key determinants of bacterial physiology and can drive phenotypic diversity. *S. pyogenes* is a fastidious pathogen with a compact ∼1.8 Mb genome and extensive nutritional auxotrophies, including requirements for multiple vitamins and most amino acids. Accordingly, it is typically grown in complex, peptide and vitamin-rich media such as Todd–Hewitt broth (THY) supplemented with yeast extract or blood. In contrast, our work examines *S. pyogenes* grown in chemically defined media that constrain nutrient availability. As a successful human pathogen, *S. pyogenes* has evolved substantial metabolic and regulatory plasticity, enabling it to adapt to fluctuating environments encountered within the host. In this context, we identified a distinct pigmented phenotype that emerged exclusively during growth in CDM that was not detectable in THY, suggesting there is suppression of metabolic or regulatory pathways under nutritionally abundant conditions. The more austere conditions of CDM have revealed specialized metabolic and signaling phenotypes like pigment development and quorum sensing regulation and may be relevant to specific host niches like mucosal surfaces or skin. Collectively, the media-dependent production of this previously uncharacterized pigment underscores the importance of environmental context in shaping bacterial traits and highlights the potential for standard cultivation conditions to overlook physiologically meaningful phenotypes.

Results indicate that oxygen is required for pigment color development, while the pigment precursor is likely synthesized independently of oxygen. Bacteria cultured anaerobically formed colorless pellets that rapidly become pigmented within minutes of exposure to oxygen. Based on this observation, we propose that oxygen facilitates a post-biosynthetic conversion that renders the pigment visible by eye. Treatment with increasing concentrations of H₂O₂ caused the pigment to appear darker, suggesting that oxidative conditions modify the pigment, potentially generating altered oxidized forms of the molecule. When anaerobically grown cells were heat-inactivated prior to oxygen exposure, the color of cell pellets remained unchanged, and argues for a need for an oxygen-dependent enzymatic step in pigment maturation, as described among bacterial pigments of carotenoid-derived compounds^18,42^. The formation of Staphyloxanthin involves a colorless intermediate that gradually acquires a yellow hue, ultimately culminating in an intense golden color upon exposure to oxygen, attributed to an enzymatic oxidative step^18,42^. Further supporting the role of oxygen, the biosynthetic gene cluster (*crtOPQMN*) responsible for pigment production in this *Staphylococcus aureus* is regulated by AirSR, a two-component system known to sense oxygen levels^43^. This underscores the critical influence of molecular oxygen in pigment maturation and the presence of an unpigmented precursor. Our findings also support that the GAS pigment precursor undergoes a redox-sensitive, oxygen-dependent enzymatic maturation step.

The heterologous expression of Staphylococcus *crtM* and *crtN* genes in *S. pyogenes* led to the production of the yellow carotenoid 4,4′-diaponeurosporene and indicates that Streptococcus can produce the precursor isoprenoid farnesyl pyrophosphate (FPP)^18^ and likely other isoprenoid intermediates. Genomic analyses reveal that *S. pyogenes* encodes enzymes of the mevalonate pathway, which is responsible for the biosynthesis of isopentenyl diphosphate and dimethylallyl diphosphate, the universal building blocks of isoprenoids^32^. However, it lacks identifiable homologs of staphyloxanthin biosynthetic enzymes, including *crtM* and *crtN* (dehydrosqualene synthase and desaturase, respectively) that catalyze condensation and desaturation of two FPP molecules, nor homologs of CrtO, P, or Q, which mediate glucosylation and fatty acid condensation of diaponeurosporenoate, suggesting that pigment production in *S. pyogenes* does not proceed via a classical carotenoid biosynthetic route^16,42^.

However, our findings do support a link between isoprenoid biosynthesis and pigment. Simvastatin, a well-known inhibitor of HMG-CoA reductase, a key enzyme in the mevalonate pathway, was used to probe this connection. Addition of statin to bacterial cultures produced a noticeable shift in pigment coloration, from a rich orange to pale yellow. Complete loss of pigmentation, however, was not observed. This phenotype suggests that isoprenoid-derived intermediates contribute to pigment chemistry, stability, or maturation, yet are unlikely to be the sole determinants of color. It remains possible that the statin, designed as a therapeutic agent to inhibit cholesterol synthesis in humans, does not fully suppress the mevalonate pathway in bacteria. Genomic analyses of streptococci do not provide evidence for a non-mevalonate isoprenoid route via 2-C-methyl-D-erythritol 4-phosphate (MEP)^32^. Thus, if an isoprenoid derivative contributes to pigmentation after full inhibition of HMG-CoA reductase, this would argue for a non-canonical biosynthetic process.

Our findings indicate that pigment production in *S. pyogenes* is tightly coordinated with purine metabolism and its associated regulatory feedback networks. Although purine biosynthesis in *S. pyogenes* has not been thoroughly investigated, insights from related organisms like *Bacillus subtills* underscore the role of interconnected feedback mechanisms in maintaining nucleotide homeostasis^44^. Under replete conditions, cells prioritize the salvage pathway over *de novo* synthesis due to lower energy requirements^45^. Inhibition of IMP dehydrogenase by MPA completely abolished pigment formation, likely reflecting disruption of guanine nucleotide synthesis and accumulation of IMP. Excess IMP is likely redirected toward AMP synthesis, leading to elevated adenine nucleotide levels and a consequent imbalance between adenine and guanine pools. Such imbalance is known to exert feedback inhibition on IMP, PRPP synthetase, and Glutamine PRPP amidotransferase, which catalyzes the first committed step of de novo purine synthesis, limiting overall purine flux. Supplementation with exogenous adenine further reinforced this repression by enhancing purine salvage, increasing intracellular AMP levels, and strengthening feedback inhibition of purine biosynthesis. Similarly, guanine supplementation inhibited pigment production in a dose-dependent manner, indicating that excess purines, whether it is guanine or adenine, are sufficient to suppress pigmentation. Together, these observations suggest that elevated nucleotide pools broadly repress purine biosynthetic activity and associated metabolic processes required for pigment formation. In contrast, removal of adenine from CDM resulted in pronounced hyperpigmentation, implying that limiting purine salvage relieves feedback inhibition and restores metabolic flux favorable for pigment synthesis. Notably, adenine depletion was partially restored pigment production in the presence of IMP dehydrogenase inhibition, underscoring the dominant role of feedback regulation rather than enzyme activity alone. However, mutants in de novo purine biosynthesis (*pur* mutants) failed to regain pigmentation under these conditions, highlighting the requirement for an intact biosynthetic pathway in addition to relief from feedback constraints. Overall, these data suggest that pigment biosynthesis in *S. pyogenes* is tightly coupled to purine metabolic homeostasis through overlapping feedback networks. When this balance is disrupted, cellular metabolism is reprogrammed away from secondary metabolite synthesis, leading to loss of pigmentation.

The repeated isolation of insertion mutations in the HAD family hydrolase *yigB* and the clean in-frame deletion of the gene that was complemented *in trans*, points to an intriguing role in pigmentation. The function of *yigB* in *S. pyogenes* has not been determined, but in *E. coli*, YigB is associated with riboflavin biosynthesis ^46^. Given that its production requires guanosine as a substrate, we speculate that either a flavin-associated metabolic pathway influences pigment production in *S. pyogenes* or that mutation in the gene leads to purine imbalance. The distinct HPLC retention time and UV-visible absorbance profile of the pigment suggests that it is unlikely to be riboflavin itself. Therefore, *yigB* may have an indirect role in modulating pigmentation through broader metabolic or redox associated pathways.

Given that a great number of transposon mutations were identified in central metabolic pathways, we are aware that a mutagenesis approach we used cannot identify essential genes under the screening conditions. Consequentially, additional genes involved in pigment biosynthesis or regulation may not have been captured in our screen. Our attempts to employ a recently developed CRISPRi strategy were surprisingly unsuccessful due the unanticipated effect that deleting the chromosomal copy of *cas9* in MGAS5005 affected pigment production. Reasons for this were not determined, but a prior report indicated dramatic regulatory and metabolic effects of deleting *cas9* in strain 5448 and points to the critical aspect of balanced central metabolic pathways^47^.

Finally, we found that the pigmented extract inhibited NFκB activity of TLR2-stimulated RAW-Blue macrophages, indicating an effect on host inflammatory signaling. NFκB is a key transcription factor that plays a central role in innate immune responses by regulating the expression of numerous inflammatory genes including cytokines and chemokines^41^. Modulation of NFκB activity by bacterial pigments through different mechanisms has been reported in several bacterial systems. For example, pyocyanin modulates NFκB activity in *P. aeruginosa* infections by increasing the production of cytokines like IL-8^48^. The pigment of *Chromobacterium violaceum* suppresses pro-inflammatory cytokines such as TNF-α and IL-6 while enhancing IL-10 production in animal models^49,50^. Staphyloxanthin and granadaene modulate NFκB through their antioxidant activity and protect the bacteria from oxidative burst within host phagocytes^18,51^. Whereas our tests found that crude and partially purified fractions of the *S. pyogenes* pigment inhibited inflammatory signaling, further purification and structure elucidation will be required to identify active components and to elucidate the underlying mechanism of action. Preliminary NMR analysis of the active fraction indicated the presence of UDP-GlcNAc and a glycopeptide, raising the possibility that the active molecule may be associated with a cell wall-related precursor or metabolite. However, the precise structure and identity of the bioactive component remains unresolved. Future work will focus on purification, structural elucidation, and mechanistic studies as necessary to determine the molecular basis of the observed immunomodulatory activity.

Taken together, results indicate that the pigment precursor in *S. pyogenes* may derive its core skeleton from either the carotenoid or purine pathway, or potentially from both. It is also possible that one pathway provides the structural backbone while the other contributes a regulatory molecule capable of modulating another pathway. Furthermore, an oxygen-dependent enzymatic step appears essential for the maturation of the precursor into a visible pigment. This study thus documents the nutrient and oxygen dependent nature of pigment production in *S. pyogenes*, while highlighting the principal biosynthetic pathways likely responsible. The differential effects of purine availability in pigment production further underscore the complexity of the regulatory networks governing pigment synthesis. Future investigations will require detailed structural characterization of the pigment and elucidation of its potential role in the observed immunomodulatory effects.

## MATERIALS AND METHODS

### Bacterial strains and culture conditions

All strains used in this work are listed in Table S1 in the supplemental material, and the construction of the various mutants and recombinant strains is described below. *S. pyogenes* strains were routinely cultured in Todd Hewitt Broth (RPI International) + 0.2% yeast extract (VWR) at 37° C with 5% CO_2_. A chemically defined medium (CDM) containing 1% glucose was used for all assays^27,52^. When necessary, antibiotics were added to the following concentrations for *S. pyogenes*: erythromycin, 0.5 µg/mL; chloramphenicol, 3 µg/mL; kanamycin, 150 µg/mL; neomycin, 150 µg/mL; spectinomycin, 150 µg/mL. *E. coli* cloning strains were routinely cultured in Luria broth (BD Difco™) with antibiotics used at the following concentrations: erythromycin, 500mg/mL; chloramphenicol, 10 µg/mL; spectinomycin, 150 µg/mL. For storage at −80° C, sterile glycerol was added to a final concentration of 20%.

**Starter cultures were prepared** by culturing single colonies of GAS in THY to an OD600 of 0.8. Glycerol was added to a final concentration of 20%, and individual aliquots were frozen and stored at −80°C. For all experiments, aliquots were thawed, and the bacteria were pelleted and resuspended in CDM before dilution into fresh CDM.

### Growth condition and visualization of a pigment

25 mL of CDM in 50-mL conical tubes was inoculated with 500 µL of a starter culture (1:40 dilution) and incubated overnight at 37°C in a 5% CO₂ incubator with caps loosely closed. The following day, cultures were shaken at 250 rpm for 30 min at 37 °C. Cells were harvested by centrifugation at 2,800 × g for 10 min at 37 °C using a JA-10 fixed-angle rotor (Thermo Scientific) and pellets were photographed to assess pigment production.

### Pigment extraction by methanol

Cell pellets of 25 mL cultures were resuspended in 1 mL of methanol containing 0.1% formic acid, then thoroughly mixed. The suspension was centrifuged at 2,800 × g for 10 min at 37 °C using a JA-10 fixed-angle rotor (Thermo Scientific) to pellet cells, and the resulting supernatant was collected and passed through a 0.22 μm syringe filter prior to downstream analysis. Spectrophotometric absorbance analysis was conducted on 1 mL of filtered extract over a wavelength range of 300 - 1100 nm, using MeOH + 0.1% formic acid as the blank.

### Pigment extraction by DMSO

Extracts were typically obtained from 50 mL cultures grown overnight in CDM in 250 mL conical Erlenmeyer flasks, harvested by centrifugation at 2,800 × g for 10 min at 37 °C using a JA-10 fixed-angle rotor (Thermo Scientific). The cell pellet was washed with PBS and centrifuged again under the same conditions. The washed pellet was then resuspended in 2 mL of DMSO containing 0.1% formic acid and mixed thoroughly. The suspension was centrifuged at 4000 rpm and 37 °C for 10 minutes to remove cellular debris. The resulting supernatant was collected and filtered through a 0.22 μm syringe filter. To 2 mL of the filtered supernatant, a mixture of 2 mL ethyl acetate and 1 mL methanol was added, and the mixture was incubated overnight at 4°C, resulting in pigment precipitation. The following day, the precipitate was centrifuged at 2,800 × g for 10 min at 37°C using a JA-10 fixed-angle rotor (Thermo Scientific). Pelleted precipitate was washed with 1 mL of ethyl acetate and centrifuged again. To determine the dry weights of the precipitate, the material was dissolved in 1 mL of Milli-Q water, transferred to pre-weighed glass vials (Fisher Scientific) and dried using a speed vac concentrator (Thermo Scientific).

### Anaerobic Culturing

25 mL CDM in 50 mL falcon tubes with lids loosely opened for gas equilibration were placed inside an anaerobic chamber 24 hours prior to inoculation. 500 μL of overnight starter cultures was used to inoculate the CDM, which was then incubated at 37 °C in the anaerobic chamber. Prior to removal from the chamber, falcon tubes were sealed with parafilm to prevent oxygen exposure during handling.

### Heat-treatment experiments

25 mL bacterial cultures were grown overnight in 50 mL Falcon tubes at 37°C + CO_2_ under static conditions (“microaerobic growth”) or in an anaerobic chamber. The following day prior to aeration, cultures were heat treated at 80°C for 30 minutes in a water bath. Cultures were then aerated for 30 minutes, followed by centrifugation for cell harvesting. The pellets was photographed to document coloration.

### pGH9:IS*S1* mutagenesis

An IS*S1* insertion library was generated by electroporating the temperature-sensitive plasmid pGh9-IS*S1* into MGAS5005 ^29^. Insertion mutants were isolated by selective plating at the non-permissive temperature (37°C), and isolated colonies were picked into 200 μL CDM cultures in the presence of 1mg/mL erythromycin in 96-well PCR tubes and covered with breathable film to expose to air. After 24 hours of incubation, cultures were pelleted and non-pigmented mutants or any displaying diminished pigment were selected. These were regrown in 5 mL CDM + Erm and phenotypes were validated. Genomic DNA of each mutant was extracted and digested with *Hind*III (1 µg DNA in a 15 µL reaction) at 37°C overnight, followed by heat inactivation at 80C for 20 min. Digested fragments were self-ligated using 1mM ATP and T4 DNA ligase at room temperature for 1 hour. Ligated products were used at template for PCR amplification using outward facing primers specific to IS*S1* element (ISS1_For_4 and ISS1_R_out_1, Table S2). PCR products were purified and sent for Sanger sequencing using primer ISS_R_out_2. Sequences were aligned with the published MGAS5005 genome to identify insertion sites.

### HPLC analysis

HPLC methods were conducted using a C18 reverse-phase column (Kinetex, 5 µm EVO C18, Phenomenex, 150 mm x 4.6 mm) with detection by DAD at 468 nm on an Agilent Technologies 1260 Infinity HPLC system (Agilent Technologies). The mobile phases used were water and methanol, each containing 0.1% formic acid. Chromatographic separation was performed using a gradient elution starting with an initial 5-minute isocratic hold at 97.5% aqueous phase (water + 0.1% FA) and 2.5% organic phase (Methanol + 0.1%FA). After equilibration, a linear gradient was applied from 5 to 25 minutes, increasing the organic phase (methanol +0.1%FA) to 100%. The column was then held at 100% organic (Methanol + 0.1%FA) from 25 to 35 minutes to elute strongly retained compounds and wash the column prior to re-equilibration.

### Construction of deletion mutants

All newly generated deletion strains in this study were derived in an MGAS5005 strain in which the restriction modification gene *hsdR* was replaced by an erythromycin resistance cassette to improve transformation efficiency^53^. The Δ*hsdR*::*ery* deletion was introduced into MGAS5005 by subcloning the region from strain NV6 (primers JC808/809^53^, Table S2) into a temperature-sensitive plasmid, pJC159 (primers JC810/811;^54^). After confirming the plasmid in *E. coli*, it was electroporated into *S. pyogenes*, and mutants were generated by selecting for allelic exchange using a two-step integration method as previously described^55^. To delete *yigB*, this same two-step, temperature-dependent method was used; primers set JC835/837, JC292/304, and JC836/838 were used to amplify the upstream flanking region, *aphA3*, and downstream flanking region, respectively, and these were cloned into pJC159 to make the knockout plasmid, pJC523. For the deletion of the other pigment-related genes, ∼1.5 to 2 kb upstream and downstream of the gene of interest and the *aphA3* kanamycin resistance gene were amplified by PCR (*fps*: upstream, JC919/920; *aphA3*, JC921/922; downstream, JC623/923; *nupPQ*: upstream, JC243/924; *aphA3*, JC925/926; downstream, JC927/928; *pfl*: upstream, SP001/JC929; *aphA3*, JC930/931; downstream, JC816/932; *atmDE*: upstream, JC346/348; *aphA3*, JC320/321; downstream, JC347/349) and assembled using the NEBuilder HiFi Assembly Kit (New England Biolabs). Microgram amounts of the linear assembled product were amplified by PCR and used to transform electrocompetent MGAS5005 Δ*hsdR*::*ery*, followed by selection on kanamycin + neomycin. Genotypes of all resulting mutants were confirmed by PCR.

### Construction of complementation plasmids

The *pfl* complementation plasmid, pJC496, was constructed by amplifying *pfl* with its native promoter by PCR (primers JC738/739) and cloning this fragment into PstI-digested pLZ12-Sp. The other complementation plasmids were constructed by cloning PCR-amplified genes downstream of a 122 bp fragment containing the *S. pyogenes recA* promoter (primers JJ92/JJ205) in pLZ12-Sp (pJC525) to make pTD01 (*yigB*, primers TD1/TD2), pJC547 (*nupPQ*, primers JC943/944), pJC548 (*atmDE*, primers JC945/946), and pJC549 (fps, primers JC947/948).

### Inhibition of protein synthesis with chloramphenicol

25 mL bacterial cultures were grown in 50 mL falcon tubes overnight at 37°C as described above, either in a CO_2_ incubator or an anaerobic chamber. Prior to aeration, chloramphenicol was added to a final concentration of 3 µg/mL, and cultures were aerated for 30 minutes. Following treatment and aeration, cells were harvested by centrifugation at 2,800 × g for 10 minutes at 37 °C using a JA-10 fixed-angle rotor (Thermo Scientific) and the color of the pellets was visualized.

### Statin treatment

Simvastatin (5 mg; Sigma-Aldrich) was dissolved in dimethyl sulfoxide (DMSO) to prepare a stock solution at 1 mg/mL. 25 ml overnight cultures were grown in the CDM in the presence of simvastatin at the indicated concentrations of 0, 3.5, 7, 14, and 28 µg/ml. Following incubation, cultures were shaken for 30 minutes on a rotary shaker. The next day, the culture was centrifuged at 4000 rpm for 10 minutes, after which pigmentation was assessed visually.

### MPA treatment

The minimum inhibitory concentration (MIC) of mycophenolic acid (AstaTech) was initially determined to be 1.25 mg/ml. Based on this result, cultures were inoculated into 25 mL of chemically defined medium (CDM) containing MPA at the sub-MIC concentration of 1 mg/ml. Cultures were incubated overnight at 37 °C, and on the following day were shaken for 30 min on a rotary shaker and centrifuged at 4000 rpm for 10 mins prior to visual assessment of pigmentation.

### Varying nucleoside concentrations

To assess the impact of exogenous nucleoside concentrations on pigmentation, CDM was prepared with variations in concentrations of adenine, guanine, or uracil. Pigment production was determined following culturing as described above.

### Disc diffusion assay

Overnight cultures of *Streptococcus pyogenes, Staphylococcus aureus,* or *Streptococcus agalactaie* grown in THY broth, or *E. coli* or *Pseudomonas aeruginosa* grown in LB broth were spread onto respective agar plates to establish lawns. After liquid dried, sterile blotting paper discs were placed onto the plates. 10 µL samples of 1 mg/ml *S. pyogenes* extracts, 3 mg/ml chloramphenicol (positive inhibitory control) or PBS (vehicle control) were placed onto filter paper. Plates were incubated overnight at 37 C.

### Total Antioxidant Capacity (TAC) assay

Total Antioxidant capacity (TAC) assay kit (Abcam) was used following the manufacturer’s instructions. Pigmented and non-pigmented extracts were resuspended in PBS at a concentration of 2 mg/mL. Trolox standards were prepared by serial dilution to generate a standard curve. For the assay, 5 µL of sample or Trolox standard was added to the wells of a clear 96-well microplate. Subsequently, 10 µL of Reagent 4 solution (peroxidase diluted in buffer at a 1:9 ratio) was added, followed by 85 µL of ABTS working solution. The plate was gently mixed and incubated at room temperature for 7 minutes to allow color development. Absorbance was measured at 414 nm using Synergy 2 BioTek plate reader. Total antioxidant capacity was calculated based on the Trolox standard curve.

### NFκB reporter assay

RAW Blue macrophages (InvivoGen), which have a chromosomally integrated NFκB-inducible secreted embryonic alkaline phosphatase (SEAP) reporter, were used to assess NFκB activation. Cells were cultured in DMEM (Gibco) supplemented with 10% heat-inactivated FBS (Gemini), penicillin/streptomycin (Corning), and 200 μg/mL zeocin (InvivoGen) in tissue culture-treated T25 flasks (Greiner BioOne). Once they reached 60–70% confluency, cells were counted and seeded into flat-bottom 96-well tissue culture plates (Corning) at a density of 2.5 × 10⁴ cells per well one day prior to the experiment. The following day, cells were stimulated with 100 ng/mL PAM3CSK4 (or left unstimulated), then treated with different concentrations of extract and incubated at 37°C + CO2 incubator. Following 18 hours of incubation, cell-free supernatants were collected. A volume of 50 μL of each supernatant was mixed with 150 μL of QUANTI-Blue substrate (InvivoGen) and incubated at 37°C + CO_2_ for 1 hour in a 96-well plate. The resulting color change was measured at 625 nm using a Synergy 2 BioTek plate reader.

### Sephadex LH20 size exclusion chromatography

Pigment was extracted from 20 liters of culture using the DMSO extraction method as described above with scaled parameters. The resulting pellet obtained after DMSO extraction was washed with ethyl acetate to remove nonpolar impurities, then resuspended in 25 mL of MilliQ water. The aqueous suspension was lyophilized to obtain dried crude extract. For chromatographic separation, the dried extract was resuspended in 8 mL of a 50:50 (v/v) methanol (MeOH): water solution. The sample was then loaded onto a Sephadex LH-20 column (400 g Sephadex, 120 cm tall and 4.6 cm internal diameter) pre-equilibrated with the same solvent. Elution was performed using a 1:1 (v/v) MeOH: water mobile phase at a constant flow rate of 0.5 mL/min. Colored bands of dark orange, orange, and yellow were apparent by eye and were collected separately for further analysis.

### Hemolysis assay

Bacterial strains were grown overnight in THY as mentioned above. The following day, cultures were diluted to an OD600 of 0.5. For spot assays, 10 µL of each suspension was placed on blood agar plates. For stab assays, a sterile inoculating loop was used to inoculate the cultures by stabbing directly into the blood agar surface. Plates were incubated at 37°C in a CO₂ incubator and hemolytic activity was assessed after 24 hours.

## ACKNOWLEDGEMENTS

We thank Natalia Korotkova for her keen observations and for providing strain 2221 and its derivatives; Debra Bessen for providing the GAS strains shown in Supplemental Table 3; Matt Henke and his lab for use of their anaerobic chamber; and Vitor Lourenzon and Alessandra Eustaquio for assistance with extract chromatography.

